# scAgeCom: a murine atlas of age-related changes in intercellular communication inferred with the package scDiffCom

**DOI:** 10.1101/2021.08.13.456238

**Authors:** Cyril Lagger, Eugen Ursu, Anaïs Equey, Roberto A. Avelar, Angela O. Pisco, Robi Tacutu, João Pedro de Magalhães

## Abstract

Dysregulation of intercellular communication is a well-established hallmark of aging. To better understand how this process contributes to the aging phenotype, we built scAgeCom, a comprehensive atlas presenting how cell-type to cell-type interactions vary with age in 23 mouse tissues. We first created an R package, scDiffCom, designed to perform differential intercellular communication analysis between two conditions of interest in any mouse or human single-cell RNA-seq dataset. The package relies on its own list of curated ligand-receptor interactions compiled from seven established studies. We applied this tool to single-cell transcriptomics data from the Tabula Muris Senis consortium and the Calico murine aging cell atlas. All the results can be accessed online, using a user-friendly, interactive web application (https://scagecom.org). The most widespread changes we observed include upregulation of immune system processes, inflammation and lipid metabolism, and downregulation of extracellular matrix organization, growth, development and angiogenesis. More specific interpretations are also provided.

## Introduction

Aging remains a poorly understood biological process despite affecting most organisms ^1^. One of the difficult aspects to model is how the dynamics of OMICs and tissue homeostasis influence each other throughout the lifespan. To gain new insights on how to bridge this gap, we focused our attention on intercellular communication (ICC). Dysregulation of ICC has been defined as a hallmark of aging ^2,3^ and has recently been proposed as one of the causes leading to the cell-to-cell stochasticity arising with age ^4^. Well-known communication deregulations include inflammaging (a chronic low-grade age-associated inflammation) ^5^, impaired immune surveillance ^6^, increase in senescence-associated secretory phenotype (SASP) ^7^, altered communication between stem cells and their niche ^8,9^, remodeling of the extracellular matrix ^10,11^ and changes in endocrine and neuronal communication ^12^. Interestingly, interventions involving extracellular signals have been shown to partially reverse some of the aging phenotypes. This includes targeting endocrine mediators such as insulin-like peptides and growth hormones ^13,14^, the use of anti-inflammatory compounds ^15–17^, heterochronic tissue transplants and heterochronic parabiosis ^18–20^.

Direct measurement of intercellular communication is complicated and usually depends on the type of mediators considered, such as surface receptors, soluble factors, extracellular vesicles ^21^ or even mitochondria ^22^. However, recent studies have shown that specific aspects of ICC can be inferred from single-cell gene expression data ^23,24^. Following the pioneer study that drafted the first comprehensive database of ligand-receptor interactions (LRIs) ^25^, and based on the development of statistical tools dedicated to building cell-type to cell-type communication networks ^26–33^, it is now becoming standard to perform intercellular communication analyses alongside the workflow of single-cell transcriptomic studies. This notably includes recent aging articles on the mouse brain ^34^, on the mouse mammary gland ^29^, on several rat tissues ^35^, on the primate cardiopulmonary system ^36^ and on human skin fibroblasts ^37^. However, the ICC analyses performed in some of those studies suffer from several limitations, as they rely on tools designed to detect interactions rather than to investigate how the interactions change between two biological conditions (e.g., young/old, healthy/sick). Indeed, the main approach so far has been to detect interactions in young and old samples independently, and then to focus on signals appearing or disappearing with age. As a result, this method does not account for interactions that are detected in both conditions but are nevertheless changing significantly, and it also disregards the magnitude of the signal variation. More importantly, this approach lacks a statistical test to assess the significance of those changes and thus to evaluate if they are due to noise or to a true biological effect.

To alleviate such limitations, we built a new statistical framework specifically designed to perform differential analysis in intercellular communication. Our resulting R package, scDiffCom, can be applied to any human or mouse scRNA-seq dataset to analyze changes in ICC between two given conditions in a given tissue. scDiffCom includes a collection of around 5000 curated ligand-receptor interactions that we have retrieved from seven publicly available resources ^26–32^. The typical output of the package is a table of detected cell-type to cell-type interactions indicating, in particular, their strength and how they are regulated between the two conditions of interest. To facilitate the interpretation of these results, we implemented an over-representation test to determine the dominant variations at the gene, cell-type or functional level. In addition, the package provides several visualization tools.

We used scDiffCom on several published scRNA-seq datasets from the Tabula Muris Senis consortium ^38^ and the Calico murine aging cell atlas ^39^ to create scAgeCom, a large-scale atlas of age-related intercellular communication changes across 23 mouse tissues. Samples obtained from male and female mice or from different experimental techniques were treated separately to avoid any confounding factors. The results are hosted and accessible *via* an online web application (https://scagecom.org) that contains both tissue-specific analyses and a global section summarizing changes shared across multiple tissues.

Our aging-related analysis supports previous knowledge regarding intercellular communication, depicting, 1) a widespread upregulation of immune system processes, inflammation and lipid metabolism, and 2) a downregulation of extracellular matrix organization, growth, development and angiogenesis. Despite significant differences across sex and experimental techniques, we were able to predict the ligands, receptors and cell types that might play key roles in such dysregulation. Due to its generality and to the large number of tissues considered, we believe that scAgeCom contains a large amount of generated data, waiting to be interpreted, which might provide the community with new therapeutic targets and new hypotheses regarding the relationship between aging and intercellular communication.

## Results

### Collection of ligand-receptor interactions from existing databases

As with other methods analyzing intercellular communication from scRNA-seq data, our approach first relied on the collection of ligand-receptor interactions (LRIs). In order to maximize the variety of interaction types, we retrieved LRIs from seven publicly available resources: CellChat ^26^, CellPhoneDB ^27^, CellTalkDB ^28^, NATMI/connectomeDB2020 ^29^, ICELLNET ^30^, NicheNet ^31^, and SingleCellSignalR ^32^ *(Methods - Fetching ligand-receptor interactions*). Several processing steps were then necessary to obtain consistent human and mouse collections of curated LRIs (*Methods - Processing ligand-receptor interactions*). This means retaining only curated interactions, converting human genes to mouse orthologs for the human-only resources, and including both simple and complex LRIs. Simple LRIs are interactions involving a single-gene ligand with a single-gene receptor, e.g., *Apoe:Ldlr*. On the other hand, a complex interaction involves heteromeric ligands or receptors, e.g., *Col3a1:Itga1-Itgb1*.

Our approach resulted in 4741 mouse LRIs (simple: 3627, complex: 1114) and 4787 human LRIs (simple: 3650, complex: 1137) directly accessible from our R package scDiffCom (see also Supplemental Data 1). Fig.1a shows how those LRIs are distributed according to their database of origin. Whenever possible, we also included in Supplemental Data 1 the sources used by each of the seven databases to curate their interactions, including references to PubMed identifiers, FANTOM5 ^25^, HPMR ^40^, HPRD ^41^, IUPHAR ^42^, Reactome ^43^ or KEGG ^44^.

**Fig.1:**
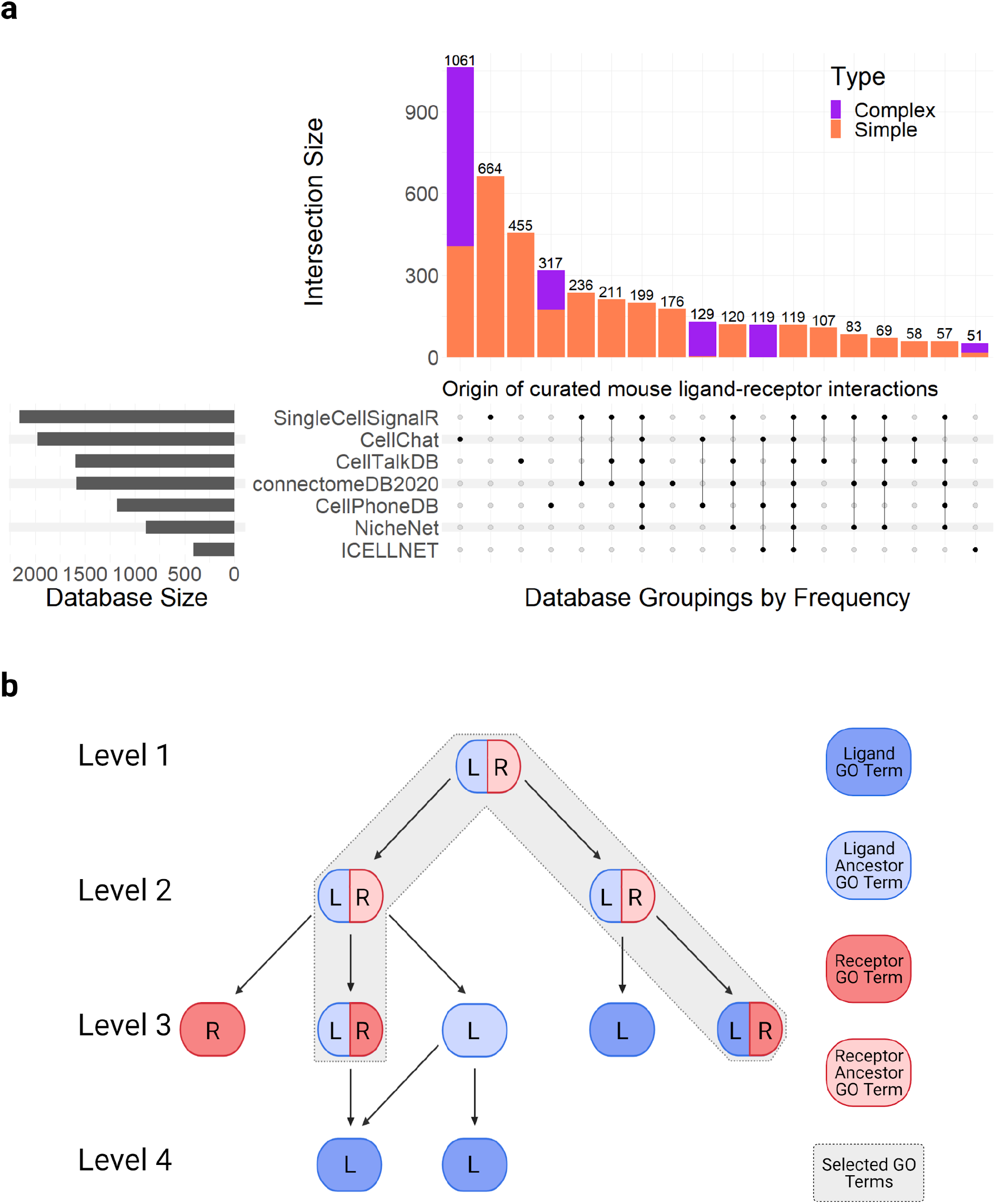
Repartition and annotation of ligand-receptor interactions in scDiffCom. a) UpSet plot showing the distribution of the 4741 curated mouse LRIs in terms of the seven databases of origin. Each column of the UpSet plot corresponds to one segment in an equivalent Venn diagram. Simplex/Complex distinguishes between LRIs composed of a single ligand gene and a single receptor gene from LRIs containing a heteromeric ligand or a heteromeric receptor. b) Representation of the method used to assign GO terms to an LRI. It consists of intersecting the nodes of the two subgraphs made from the GO terms and ancestor GO terms of the ligand (blue) and receptor (red).

### Functional annotation of ligand-receptor interactions

We annotated all LRIs with a standardized and consistent framework (*Methods - Annotating ligand-receptor interactions with GO terms and KEGG pathways*). We first associated gene ontology (GO) terms ^45^ to each interaction in a way that conveys biological meaning related as much as possible to the interaction itself rather than to each gene independently. Simply taking the intersection between the ligand GO terms and receptor GO terms would have resulted in a significant number of empty intersections (as most genes are annotated with specific GO terms and not all parent terms). Instead, as shown in Fig.1b, we associated to each LRI all GO terms formed by the intersection of two sets of nodes: 1) the nodes of the GO graph made of the ligand GO terms with their ancestors and 2) the nodes of the corresponding receptor GO graph. Since this method is prone to attaching lowly informative terms (namely those near the root of the GO graph), we also computed and indicated the level of each GO term (namely its depth in the GO graph) to facilitate downstream analysis. In addition to GO terms, we added KEGG pathways ^44^ to each LRI if all genes present in the interaction were part of the pathway. All annotations are directly accessible from scDiffCom.

### Differential cell-cell communication analysis with scDiffCom

We designed a bioinformatics method, available within the R package scDiffCom, to detect cell-type to cell-type communication patterns that significantly change between two conditions in a given scRNA-seq dataset (Fig.2). Prior to the analysis, the dataset must be formatted as an R Seurat object ^46–48^ and contain cells annotated by cell types and by the two conditions on which the differential analysis will be performed. The package then considers all possible interactions between a cell-type pair, based on the ligand-receptor interactions described in the previous section. We call each of those potential signals a cell-cell interaction (CCI). Each CCI is then assigned a score (independently in each condition) based on the expression of the genes in the respective cells (*Methods - CCI score based on the geometric mean*). A statistical procedure comprising two tests is finally performed to establish how likely each CCI is to correspond to an actual biological interaction and to be significantly differentially expressed between the two given conditions.

**Fig.2:**
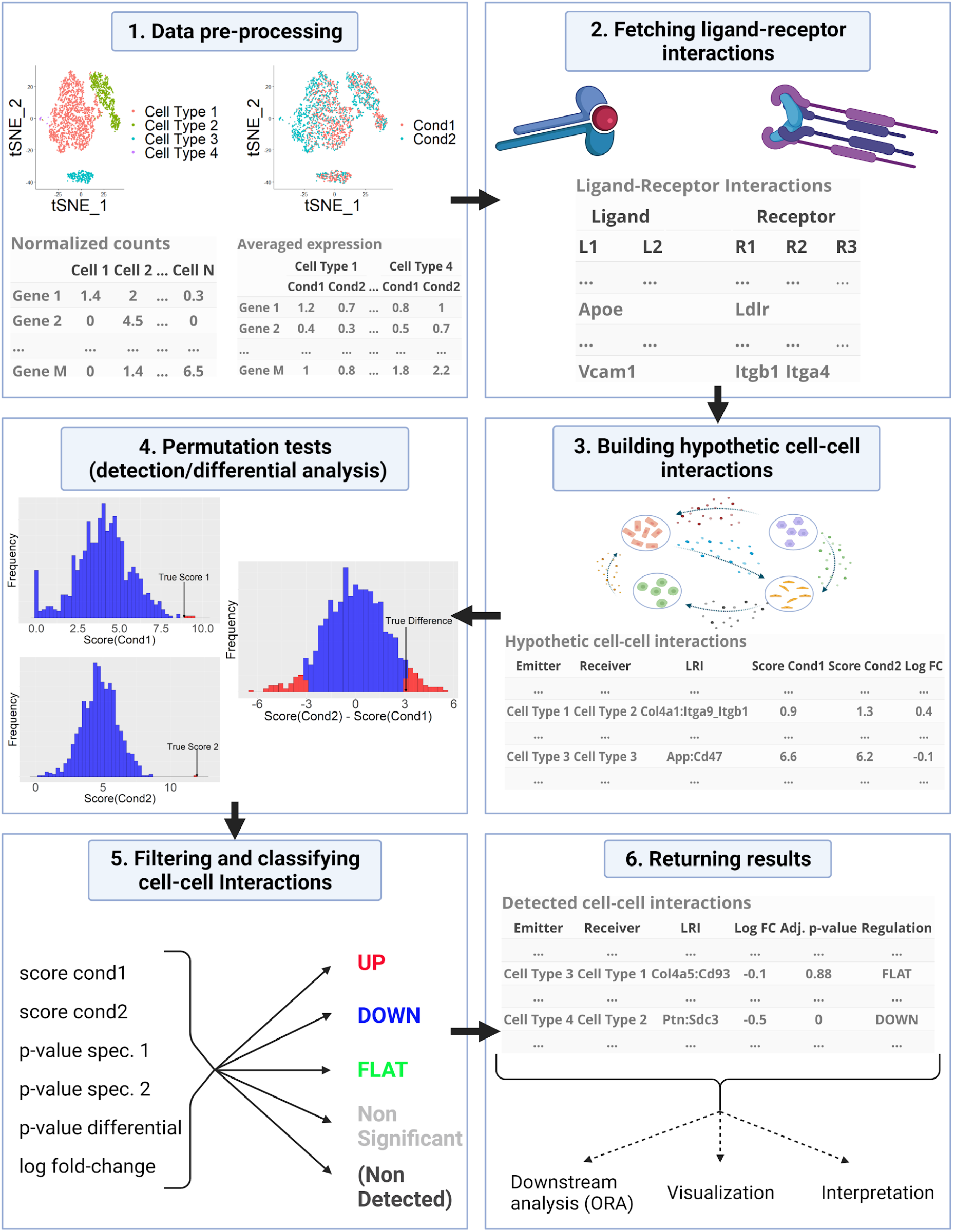
Workflow summary of scDiffCom. Read counts/UMIs from the single-cell dataset are aggregated by cell types and conditions (1). Genes are then joined with our database of ligand-receptor interactions (2) to build all the potential cell-cell interactions that can occur between cell types (3). Statistical permutation tests are then performed to evaluate the biological significance of each CCI and its differential expression (4). They are then classified based on several computed variables such as their scores, p-values and log fold-change (5). Results are returned in a convenient format for downstream analyses and interpretation (6).

The first test consists in assessing the biological relevance of each CCI (in each condition independently) based on combining previously published approaches (*Methods - CCI detection and differential analyses*). To be considered detected, a CCI has to: 1) not be lowly expressed (based on the number and percentage of cells expressing each gene), 2) be *specific*, as originally defined by the authors of CellPhoneDB ^27,49^ (this relies on a permutation test explained below), and 3) have a large enough score (compared to the other remaining CCIs). The permutation test shuffles the cell-type annotation attached to each cell to estimate how specific to a given emitter-receiver cell-type pair a particular ligand-receptor interaction is. This allows the removal of non-specific CCIs that are likely not biologically relevant.

Alongside the detection analysis, we implemented a second permutation test to assess if the score of each CCI significantly changes between the two conditions of interest *(Methods - CCI detection and differential analyses*). This consists of randomly exchanging the condition label of each cell to see whether the test statistic, the difference between the CCI scores of each condition, is different from zero. Choosing permutation tests was motivated by the fact that they are non-parametric, i.e. they make no assumptions regarding the distributions of the underlying variables ^50^. This was particularly useful in this context, as there is no obvious way to model the CCI score distribution. In addition, permutation tests can be applied to unbalanced and low sample size scenarios, which frequently appear in single-cell studies.

Finally, based on both the detection and differential analysis, each CCI is classified into one of four possible categories, further referred to as “regulation” (*Methods - CCI classification*). It can either be up-regulated (UP), down-regulated (DOWN), stable (FLAT), or correspond to a non-significant change (NSC). As such, the main output of scDiffCom is a table that contains all detected CCIs with relevant information including their regulation, log fold-changes, adjusted p-values and scores for each condition.

We note that the most computationally demanding part of this workflow is to perform the permutation tests and therefore special care was taken to optimize them. Moreover, scDiffCom can easily be run in parallel. A toy model analysis (1000 permutations on a dataset of 1000 cells and 5 cell types) takes a couple of minutes if run sequentially on a single-core computer. A more realistic example (10000 permutations on 3107 cells and 16 cell types) was measured to take around 9 minutes when run in parallel on 30 cores.

### Over-representation analysis and visualization

Every single detected CCI returned by our analysis corresponds to a communication signal whose change (or absence of change) between the two conditions can be interpreted on its own. However, as it is typical to detect several thousands of CCIs in a single tissue, scDiffCom also performs an over-representation analysis (ORA) to extract the dominant differential patterns (*Methods - Fisher’s exact test to find over-represented signals*). ORA measures the statistical association between CCI classes (UP, DOWN or FLAT) and CCI features of interest, e.g., gene, cell-type or functional annotation. For instance, it can evaluate if a given ligand-receptor interaction takes part in up-regulated CCIs more than expected by chance. Results can be sorted according to an over-representation score that combines the odd ratio (OR) and adjusted p-value returned by the test: *ORA score* = *log*_2_ *OR* · (− *log*_10_ *adj. pval*).

ORA is performed on the following categories: LRIs, ligands, receptors, emitter-receiver cell-type pairs, emitter cell types, receiver cell types, GO terms and KEGG pathways. It is useful to analyze ligands and receptors alone to see if some ligands take part in various regulated CCIs involving different receptors (or vice-versa). The same logic is valid for the cell types. Moreover, scDiffCom offers the opportunity to perform ORA on additional user-defined attributes.

Finally, scDiffCom provides two tools to produce images of the over-represented results (*Methods - Visualization Tools*). First, the function PlotORA displays the top over-represented signals of a given category and regulation with their odds ratios and adjusted p-values. Second, the function BuildNetwork creates and displays a network representation of the over-represented cell-type pairs, as will be illustrated below. Other types of data visualization can be created from direct calls to standard R graphic packages and are therefore not implemented in scDiffCom.

### Preparation and annotation of scRNA-seq aging datasets

We used the developed package to create a comprehensive murine atlas of age-related intercellular communication changes by applying scDiffCom to scRNA-seq datasets obtained from the Tabula Muris Senis (TMS) consortium ^38^ and the Calico murine aging cell atlas ^39^ *(Methods - Fetching and preparing scRNA-seq datasets*). These studies offer high-quality cell-type annotations: following standard cell clustering techniques ^51^, TMS relied on experts to manually curate their cell-type classification, whereas Calico assigned cell types to their cells based on a semi-supervised neural network using the first Tabula Muris study ^52^ as training data.

After data pre-processing, we retained 58 distinct datasets corresponding to 23 organs distributed across 5 categories: *TMS FACS (male), TMS FACS (female), TMS Droplet (male), TMS Droplet (female) and Calico Droplet (mal*e) (Fig.3). *FACS* and *Droplet* refer to the two experimental techniques used by the original studies. We kept male and female samples separate to avoid possible confounding effects. Cells from different time points were classified as either young or old, as the two conditions on which scDiffCom performs the differential analysis. To integrate the 5 types of datasets and facilitate comparisons, we standardized the cell-type names across datasets, regrouped very specific categories into broader ones, and annotated them with cell-type families (*Methods – Cell-type characterization*).

**Fig.3:**
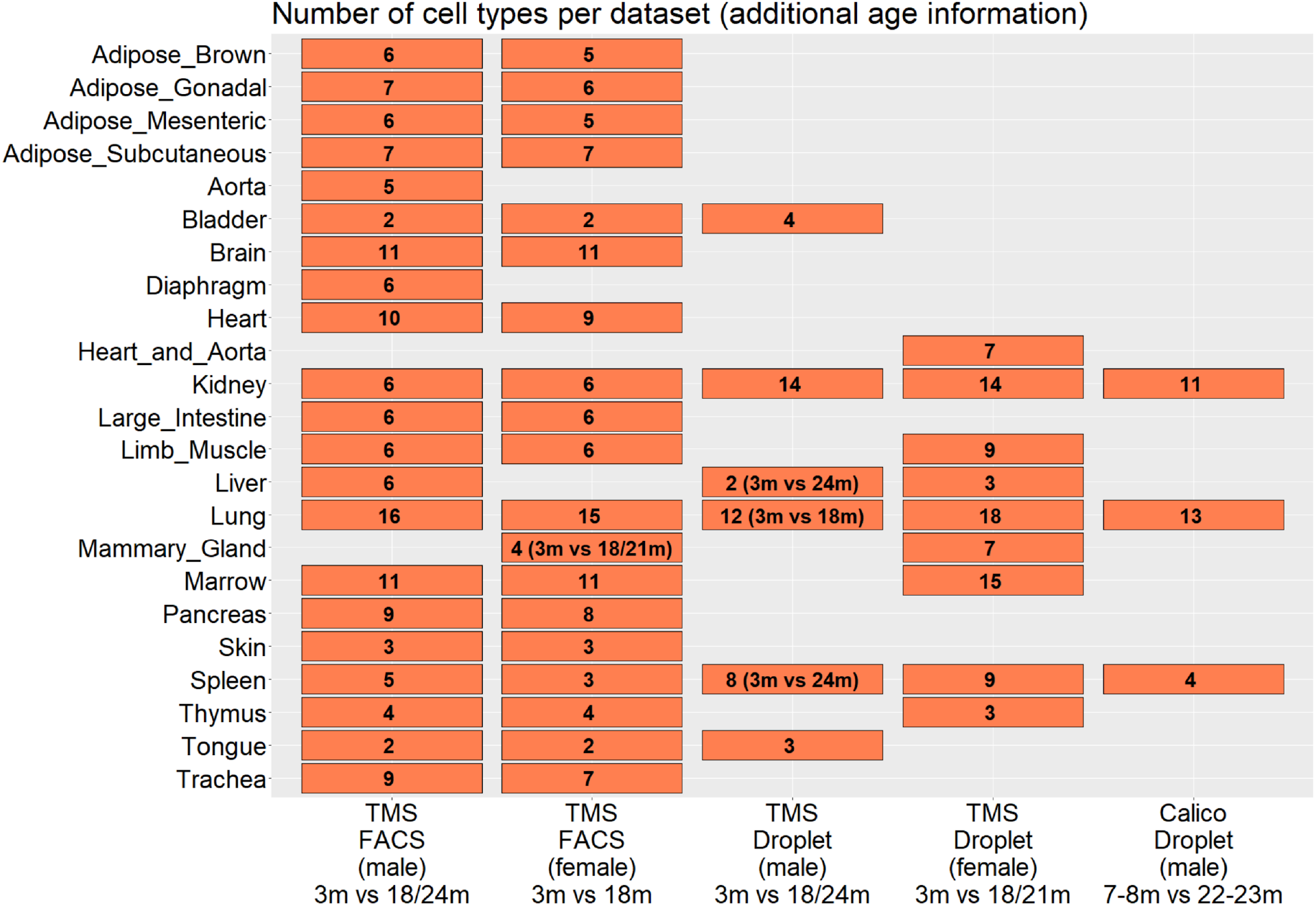
The 58 aging scRNA-seq datasets used to build scAgeCom. Repartition of the datasets according to their experimental/sex category (x-axis) and the tissue concerned (y-axis). Each rectangle represents a dataset and contains the number of detected cell types. The x-axis labels show information about the age of the mouse samples being compared.

The dataset preparation revealed that most tissues do not share the same cell types across the five experimental and sex categories. As shown in Fig.3, the number of cell types in the lung varies from 12 to 18. Moreover, some important cell types were not captured at all due to technical limitations (e.g., adipocytes in adipose tissues). As such, one must keep in mind that our results can only convey a partial representation of what is expected to happen *in vivo*.

### scAgeCom: a mouse aging atlas of intercellular communication

We applied scDiffCom to each of the 58 aforementioned datasets, performing differential analysis between young and old cells using 10000 permutations and default parameters (as defined in *Methods*). To facilitate access to this large amount of data, we created a Shiny application (*Method - Building and deploying scAgeComShiny*). This website, available at https://scagecom.org, provides both a tissue-specific section, displaying results for each dataset independently and a global section summarizing the dominant signals shared across datasets.

Fig.4a-c illustrate the interactive tables and plots provided to explore detected CCIs and their age-regulation in each dataset. Several filtering options are available to restrict interactions to a subset of cell types, LRIs or functional annotations. Results of the over-representation analysis are available as graphs and tables for cell-type-centric, gene-centric and function-centric investigations (Fig.4d-f).

**Fig.4:**
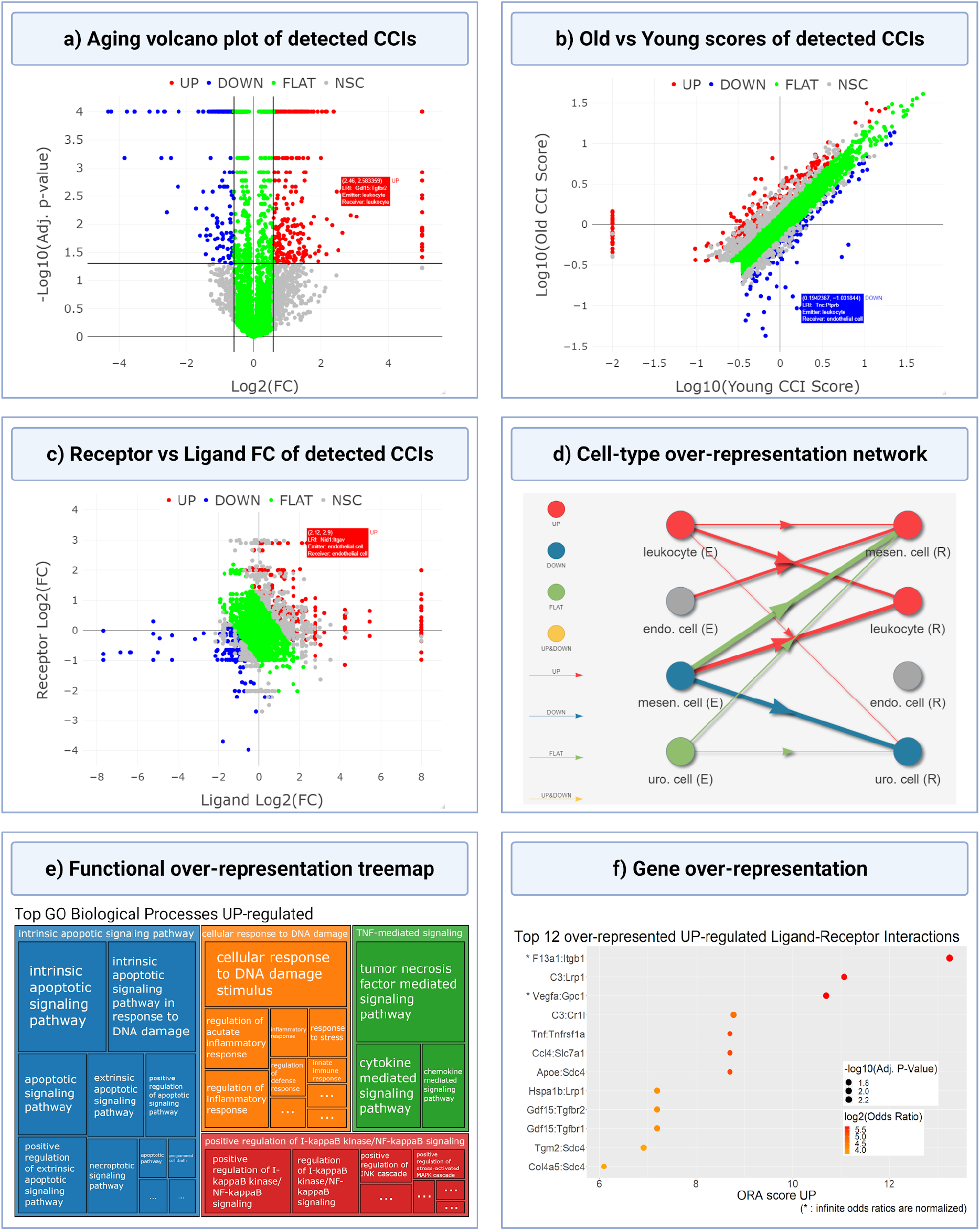
Visualization of tissue-specific analysis results in scAgeCom. Those plots are available for each of the 58 datasets present in our atlas scAgeCom. As an illustration, they are given here for the *Bladder* from *TMS Droplet (male)*. a) The volcano plot shows the CCI distribution by aging log fold-change and adjusted p-value and highlights the differences among the four classes of CCIs. b) The score plot displays CCIs in terms of young and old scores, allowing for better visualization of the initial and final values that yield the absolute CCI score changes with age. For example, it can discriminate between CCIs undergoing similar change with age but having different young and old scores. This plot also highlights stable (FLAT) CCI with high scores, which may play an important role in intercellular communication regardless of age. c) The Ligand vs Receptor FC plot captures the log fold-change of the ligand and the log fold-change of the receptor. Our approach combines the ligand and receptor expressions into a single score, thereby losing some information, which this graph recovers *a posteriori*. It illustrates if the change with age in a given CCI is driven by the ligand or the receptor. d) Network representing which emitter cell types, receiver cell types and cell-type pairs are over-represented as either up-regulated, down-regulated, stable, or both up and down-regulated with age. The latter scenario is realized when both up and down-regulated LRIs are significantly present. e) GO Treemap is obtained by merging over-represented GO terms by semantic similarity and provides a useful high-level summary. Here, only up-regulated biological processes are shown, but the same plots are available for down-regulated terms as well as GO molecular functions and cellular components. f) Top over-represented LRIs, based on their ORA score. Here we show up-regulated interactions, but similar plots are available for the other cases, as well as for ligand and receptor genes alone. Marker size and color encode adjusted p-value and odds ratio respectively, which together determine the ORA Score.

The high number of datasets analyzed allowed us to perform a global analysis to investigate age-related communication changes shared across multiple tissues. We used the signals returned by the over-representation analysis in each dataset. For each annotation, e.g., the GO term *T cell differentiation* or the LRI *B2m:Cd3g*, we counted the number of datasets in which it was over-represented (Fig.5). This allowed us to create tables with the keywords that were overrepresented in the largest number of tissues.

**Fig.5:**
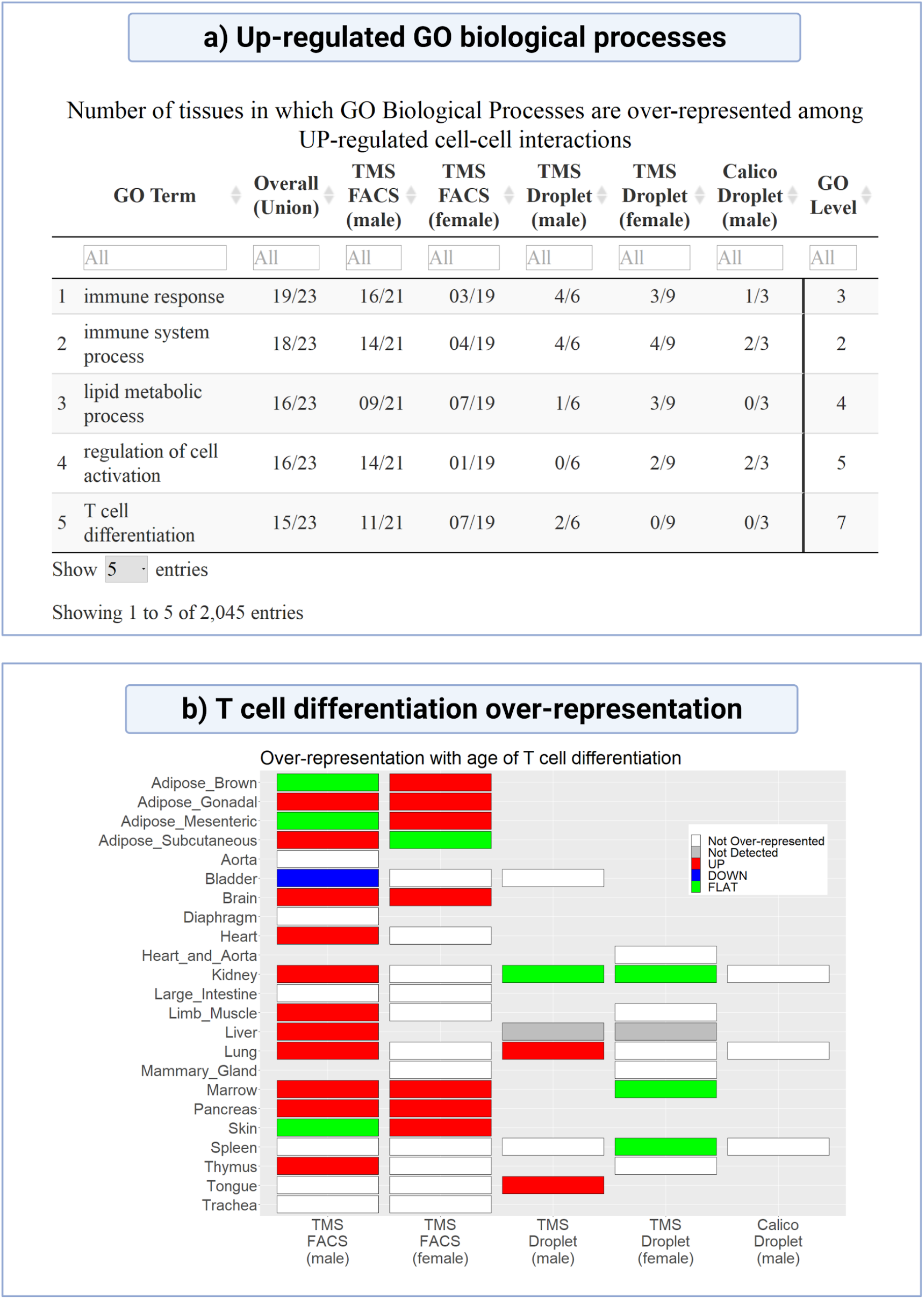
Presentation of the cross-tissue results available in scAgeCom. a) For each category on which over-representation analysis has been performed and for each regulation with age (UP, DOWN, FLAT), we provide a table that summarizes the number of tissues in which a particular keyword is over-represented. An example is given here for the up-regulated GO biological processes. b) For each keyword of any category, we also provide a summary plot that shows how it is over-represented across the 58 datasets we analyzed. An example is given here for the GO term *T cell differentiation*.

### Aging dysregulates multiple aspects of intercellular communication

Over the 58 datasets considered, we detected 382522 CCIs (5% UP, 13% DOWN, 56% FLAT, 26% NSC), corresponding to an average of 98 detected LRIs (SD = 77) between any cell-type pair. The nature and age-regulation of the detected CCIs strongly vary across datasets, sex and experimental techniques, as shown in Fig. 6. The tissues from *TMS FACS (male)* clearly show a larger fraction of down-regulated CCIs compared to all other datasets including those from *TMS FACS (female)*. Unsurprisingly, we also observe more variability and noise (namely more NSC CCIs) in *FACS* compared to *Droplet*. Those observations are partially explained in the Discussion below.

**Fig.6:**
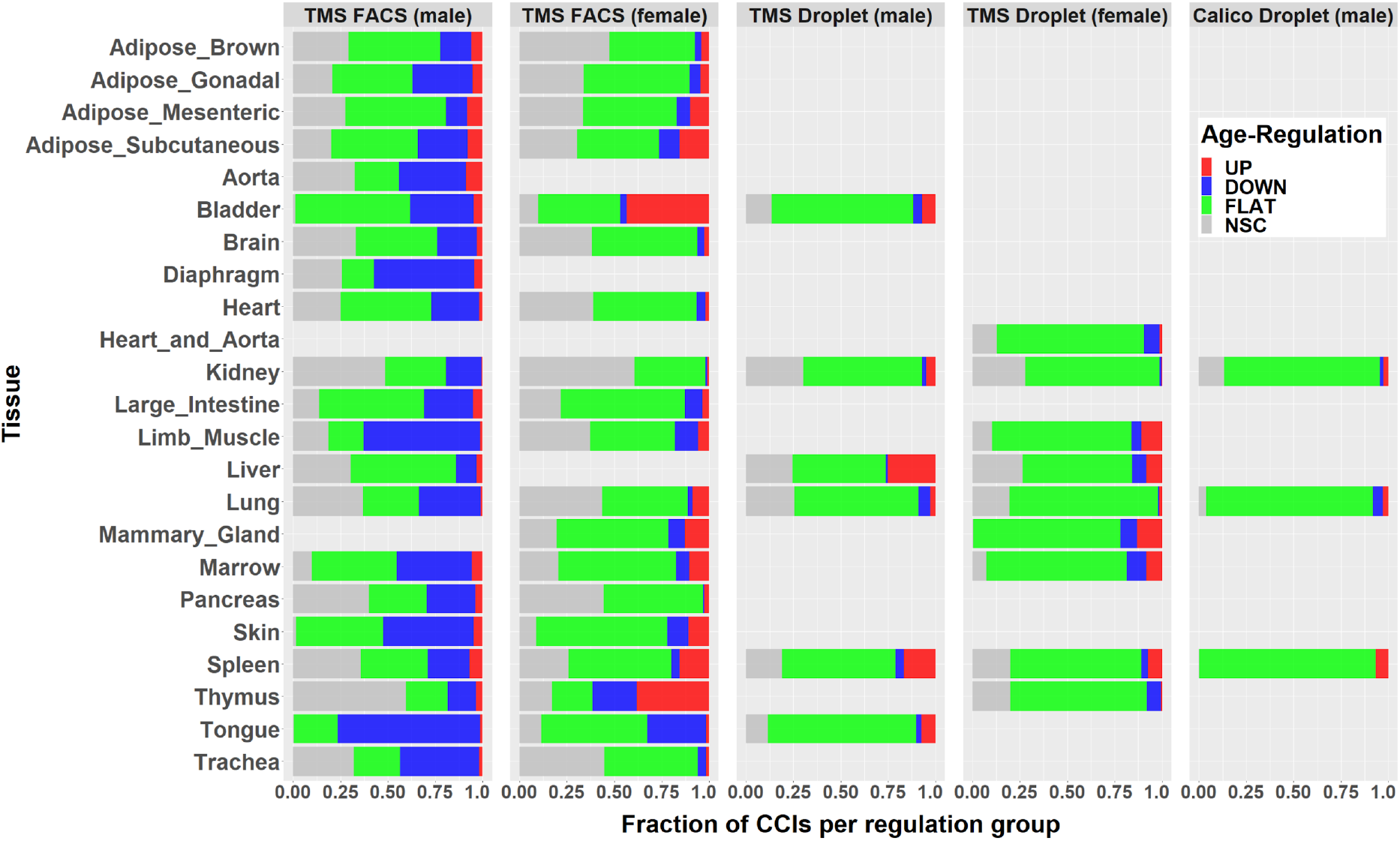
Stacked bar representation of the age-regulation of CCIs across tissues and data source. For each of the 58 datasets, the colored bar indicates the percentage of UP, DOWN, FLAT and NSC CCIs.

The global section of the atlas reveals several broad mechanisms dysregulated with age. First, consistently with the literature, we observe a major up-regulation of inflammatory and immune system processes. GO terms over-represented in more than half of the tissues include *immune response, immune system process, defense response, T cell differentiation, cytokine/chemokine-mediated signaling pathway, lymphocyte migration, positive regulation of adaptive immune response, regulation of B cell activation, regulation of viral life cycle/process* and KEGG pathways such as *antigen processing and presentation, cytokine-cytokine receptor interaction, Epstein-Barr virus infection*. Analysis of cell-type families reveals that the interaction *leukocyte-leukocyte* is up-regulated in 16 tissues and that the interactions from *endothelial, connective* and *stem* cells towards *leukocytes* are each up-regulated in at least 5 tissues. From the gene perspective, the interactions that are overrepresented in the highest number of tissues (9 to 15 out of 23 tissues) include *B2m:Cd3g, B2m:Cd3d, Tnfsf12:Tnfrsf12a, H2-D1/K1/Q6:Cd8b1, Mif:Cd74, Hmgb1:Thbd*, and *Ccl5* interacting with different chemokine receptors such as *Ccrl2, Ccr1 and Ccr5* in 6, 4 and 4 tissues, respectively.

A second up-regulated process is related to lipid metabolism, with GO terms like *lipid metabolic process, cellular lipid metabolic process, lipid catabolic process, lipoprotein metabolic process, regulation of lipid localization, fatty acid biosynthetic process. Apoe* is over-represented as an up-regulated ligand in 16 tissues, mostly via its interactions with *Sdc4, Lrp1* and *Ldlr*. However, this pattern is mainly found in *TMS FACS (male)* and happens to be reversed in four *TMS FACS (female)* tissues. Along the same line, *App:Lrp10*, is over-represented among down-regulated signals in 8 tissues from *TMS FACS (male)* but up-regulated in 4 tissues from *TMS FACS (female)* and 3 tissues from *TMS Droplet (female)*.

The most over-represented processes among signals down-regulated with age are related to the extracellular matrix (ECM) and adhesion. This includes the GO terms *biological adhesion, cell adhesion, extracellular matrix organization, extracellular structure organization, cell junction assembly, cell-matrix adhesion* and the KEGG pathways *ECM-receptor interaction, focal adhesion, adherens junction*. The main corresponding LRIs involve collagens, fibronectins and laminins interacting with integrins, mostly between connective tissue cells, epithelial cells and endothelial cells.

In addition to the changes undergone by the ECM, interactions related to growth, development, survival, differentiation and angiogenesis are also significantly down-regulated with age. Relevant GO terms include *anatomical structure morphogenesis/development, cell morphogenesis, developmental process and growth, response to growth factor, regulation of cell cycle, Notch signaling pathway, regulation of angiogenesis* and KEGG pathways include *PI3K-Akt signaling pathway, Wnt signalling pathway*. Genes such as *Fgf1, Wnt5a, Bmp2, Angpt1, Tgfb1, Tgfb3, Vegfa, Bmp6, Notch2, Fgfr1, Fgfr2, Tgfbr2, Egfr, Gpi1* are over-represented in down-regulated CCIs in 5 to 13 tissues. Cell-type pair over-representation also indicates down-regulation of communication among stem cells (6 tissues) and from stem cells towards endothelial cells (5 tissues).

The results reported so far only represent a fraction of the global signals present in our atlas. Tissue-specific results provide even more leads to be investigated, including age-regulated CCIs specific to an organ or cell type and therefore not showing up as over-represented. For instance, we noticed the down-regulation of multiple communication links towards intestinal crypt cells and *Lama2* in the large intestine; the up-regulation of *Sost* (sclerostin) and communication links towards podocytes in the kidney; the down-regulation of *Notch signalling* in the bone marrow; and the down-regulation of *Hgf* and *Tgf1b* in hepatic sinusoidal endothelial cells. Unfortunately, investigating all these signals is outside the scope of this article; however, we hope that scAgeCom will encourage other researchers to access our database for this purpose.

## Discussion

In this work, we present a package to perform differential intercellular communication analysis as well as a comprehensive database of age-related mouse cell-cell interactions. From a technical point of view, our results illustrate the importance of performing a proper statistical analysis when comparing intercellular signals extracted from scRNA-seq data. Variability and noise were indeed responsible for the classification of 26% of the detected CCIs as NSC interactions (*non-significant change*), namely those with a fold change larger than 1.5 but a non-significant adjusted p-value. Had we not used a statistical test and based our analysis solely on the appearance and disappearance of CCIs between the two conditions, as in some previous studies, 82.5% of those NSC signals (i.e., 82256 CCIs) would have been falsely considered to be age-regulated. Moreover, such an approach would have missed all CCIs detected in both conditions but nevertheless showing a significant change with age, namely 16678 interactions in our case.

Fig.6 shows that there are significant disparities in the results depending on experimental techniques and sex. Previous comparisons of single-cell sequencing techniques ^53^ have claimed that Drop-seq, by using unique molecular identifiers (UMI), is less subject to amplification noise than Smart-seq2, potentially explaining why we observe less NSC CCIs in *TMS Droplet* and *Calico Droplet* than in *TMS FACS*. The other differences can be explained by several factors. First, different datasets sometimes compare different age groups (Fig.3). Second, due to experimental limitations, captured cell types are rarely the same between two datasets of the same tissue. For example, as previously shown in Fig.3 for the Lung, there are 16 cell types in *TMS FACS (male)* against 12 cell types in *TMS Droplet (male)*, resulting in more detected CCIs in the former than the latter dataset (22540 *vs*. 11372) and in a different distribution of the percentage of UP/DOWN/FLAT/NCS CCIs (Fig.6). We also emphasize that the pronounced down-regulation observed in *TMS FACS (male)* datasets (Fig.6) was reported in a previous study performed on TMS datasets ^54^.

We now provide more specific interpretations of the main biological results reported above. Regarding the increase of immune response processes with age, we observed that the ligand *B2m* (β2-microglobulin) is over-represented among up-regulated CCIs in 17 tissues. *B2m* has already been recognized as a gene consistently overexpressed with age ^55^ and as a pro-aging circulating factor whose elevated level negatively affects cognitive functions and neurogenesis in the mouse hippocampus ^56^. Our results indicate that the increased secretion of the ligand *B2m* with age is systemic and appears to target T cells *via* their receptors *Cd3g, Cd3d* and to a lesser extent *Cd247* (explaining the occurrence of the GO term *T cell differentiation*). This could be a sign of increased antigen presentation on MHC class I. Such a widespread pattern might also indicate detrimental effects of *B2m* not limited to the brain and reinforce the idea that this protein might be a potential therapeutic target, as previously suggested ^57^.

Regarding the increase in inflammation with age, our results show a global over-representation of the terms *cytokine-* and *chemokine-mediated pathways*, and of the ligands *Ccl5, Mif* and *Hmgb1*. Those ligands are known Senescence-Associated Secretory Phenotype (SASP) factors, but we cannot confirm from our analysis whether they are really emitted by senescent cells. Follow-up studies, such as for example a cross-analysis between scAgeCom and the SASP atlas ^58^, might allow us to explore such senescence-related hypotheses more precisely. Here, we only mention that the two most over-represented up-regulated LRIs that we reported earlier, *Mif:Cd74* and *Hmgb1:Thbd*, actually seem to play compensatory roles in the context of senescence and inflammation. Indeed, it has been reported that macrophage migration inhibitory factor (Mif), a pleiotropic cytokine, can prevent cellular senescence and rejuvenate mesenchymal stem cells from age-induced senescence via CD74/AMPK/FOXO3a and autophagy in both rats and humans ^59,60^. Along the same line, the high-mobility group box chromosomal protein 1 (Hmgb1) has proinflammatory effects when binding to RAGE/Ager ^61^ but its sequestration by thrombomodulin acts as an anti-inflammatory mechanism ^62^. According to our results, *Hmgb1:Thbd* is over-represented as up-regulated in 9 tissues and *Hmgb1:Ager* in 3 tissues, pointing towards a global over-emission of *Hmgb1* with age that tissues might try to compensate by over-expressing *Thbd*.

Changes in lipid metabolism with age are known to have an important impact on the lifespan and age-related diseases ^63^. Our most intriguing results concern the widespread deregulation of *Apoe*, its receptor and *App*. We indeed observed a general over-emission of *Apoe* and an under-emission of *App* in most tissues at least in *TMS FACS (male)*. In the brain, those genes are known to play a role in the pathology of Alzheimer’s disease (AD). *App* is the precursor of amyloid beta (Aβ) peptides ^64^, *Apoe* is a known regulator of Aβ clearance ^65^ and interactions between *App* and *Apoe* receptors influence Aβ metabolism and toxicity ^66,67^. Much less research has been performed on the function of *App* outside the central nervous system ^68^, but some studies point towards its role in several pathologies such as in obesity ^69,70^, in the skin ^71^, in the intestine ^72^, where it is modulated by diet ^73^, and in muscles ^74^, particularly at neuromuscular junctions ^75^. Taken together, those studies and our results suggest that changes with age in *App* intercellular trafficking might lead to a variety of tissue-specific diseases.

The two main down-regulated processes reported in our results, namely changes in the ECM and development/growth/proliferation, are partially interdependent. They have in common the under-expression of integrins, which typically act as bidirectional mediators between the cytoplasm and the extracellular space and regulate mechanisms such as cell migration, adhesion, proliferation, apoptosis, tumor progression and senescence ^76^. The multiple functions of these proteins and their deregulation that we observe across multiple organs (most notably when they interact with collagen) suggest they might play key roles in the structural decline of tissues with age. They are also consistently down-regulated in signaling involving stem cells, potentially impacting their maintenance and homeostasis, as suggested in previous articles ^10^.

A subpart of the development/growth/proliferation pathway concerns the global down-regulation of angiogenesis and qualitative changes occurring in vessels. Several studies have previously reported a decline with age in capillary density and in the formation of new blood vessels ^77,78^, even leading some authors to postulate an “angiogenesis hypothesis of aging” ^79^. Our analysis reveals that the pair *Gpi1:Amfr*, mainly detected from or to endothelial cells, is over-represented among down-regulated CCIs in 11 tissues. Glucose-6-phosphate isomerase functions as an autocrine motility factor that stimulates endothelial cell motility ^80^. This LRI could be an important regulator of microvascular aging and its therapeutic potential seems worth investigating. Moreover, we found down-regulation of angiopoietin-1 (*Angpt1*) in 7 tissues, which could lead to the formation of leaky vessels, as vessel stability relies on the balance of Angpt1 and Angpt2 ^81^.

Finally, we conclude with some of the limitations of using gene expression of ligands and receptors to infer intercellular communication activity. First, using mRNA counts as a proxy for the actual level of secreted proteins may lead to an overestimation of some intercellular interactions, as we assume that all transcripts participate in signalling, even though a fraction of them might be produced for intracellular processes. Second, it is not always clear how some interactions (e.g., *App:Lrp10*) are shared between cells, and which fraction of them are autocrine rather than paracrine. Third, by only looking at ligands and receptors, our method does not assess changes in other players acting in downstream signalling. Fourth, we do not consider all possible types of intercellular mediators and we do not investigate inter-tissue interactions such as endocrine signals. Despite these limitations, we hope that our atlas will be useful for the community and lead to new hypotheses on intercellular communication and aging to be further tested.

## Methods

### Fetching ligand-receptor interactions

We downloaded LRIs from the most recent versions (as of March 21, 2021) of seven publicly available databases. Data from CellChat, NicheNet, and SingleCellSignalR were directly accessed from their associated R packages. Data from CellPhoneDB, CellTalkDB, connectomeDB2020 and ICELLNET were fetched online from their respective websites. All details regarding download dates and links are directly accessible from our package scDiffCom. We initially also retrieved LRIs from an eighth database, LRBaseDb from scTensor ^33^, but we did not consider it further as all curated LRIs present in it were derived from the seven others mentioned above.

### Processing ligand-receptor interactions

We analyzed the documentation and annotations of each resource to only keep their curated LRIs. We removed the interactions that were only bioinformatically predicted, such as from protein-protein interaction networks. We checked that gene symbols were HGNC approved ^82^ and, when mouse data were available, MGI approved ^83^. If mouse LRIs were not provided (namely for CellPhoneDB, connectomeDB2020, ICELLNET, NicheNet and SingleCellSignalR), we converted human LRIs to their mouse equivalent by retrieving high confidence orthology information from Ensembl version 102 ^84^ accessed through the R package biomaRt ^85^.

LRIs from each resource were combined in a single list. Special care was taken to avoid duplicates arising from the same interactions given in opposite directions (e.g., G1:G2 versus G2:G1), typically for juxtacrine signalling where the notion of ligand or receptor can sometimes be arbitrary.

As some of the resources only provide simple interactions, some of their LRIs could be incomplete. To partially correct this effect, we removed simple LRIs present in such databases if they were also found in complex databases, but only in a complex form. For instance, we removed Col3a1:Itgb1, given in SingleCellSignalR, as it always appears in a complex form in CellPhoneDB, such as in Col3a1:Itga1-Itgb1.

We manually verified the combined list of LRIs and removed about 200 of them that we considered miss-curated, e.g., *Mapk1:Fgfr2, Calm1:Adcy8, Gnas:Adcy1, Hsp90aa1:Cftr*. To do so, we first annotated each gene with descriptions from *MyGene*.*Info* ^86,87^ and categories from *OmniPath* ^88^. We then identified genes that seemed unlikely to participate in intercellular communication. Finally, we explored the interactions involving those genes and evaluated the evidence supporting their existence. Some of the removed LRIs relied on publications describing intracellular interactions that had been misinterpreted as intercellular, e.g., Loo et al. ^89^ is presented as evidence for *Hsp90aa1*:*Cftr*, but the paper does not mention intercellular communication.

### Annotating ligand-receptor interactions with GO terms and KEGG pathways

LRIs were annotated with GO terms by following the graph-based approach in Fig.1b. GO terms associated with each gene were retrieved from Ensembl version 102 ^84^ accessed through the R package biomaRt ^85^. For a given LRI, we used the R package ontoProc ^90^ to build the ligand and receptor ontology subgraphs, whose nodes included their respective GO terms and ancestors up to the root node. For complex LRIs including, e.g., multiple ligand genes, we considered the union of the terms associated with each ligand gene. The final LRI GO terms were those present in both the ligand and the receptor subgraphs, i.e., the intersection of those graphs at node-level.

To annotate LRIs with KEGG pathways, we used the R package KEGGREST ^91^ to retrieve all pathways associated with a given gene. For each LRI, we then only retained the pathways that included both the ligand gene(s) and receptor gene(s).

### CCI score based on the geometric mean

For each cell-cell interaction in the form (emitter cell type, receiver cell type, ligand(s), receptor(s)), a score φ is computed in each condition as the geometric mean 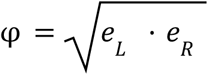 between the averaged expression *e*_*L*_ of the ligand gene in the emitter cells and the averaged expression *e*_*R*_ of the receptor gene in the receiver cells (based on normalized non-log-transformed read counts/UMIs). In the case of complex LRIs with multiple ligand genes (or receptor genes) involved, *e*_*L*_ (or *e*_*R*_) is given by the minimum value from the set of average expressions of those genes.

Defining a CCI score is a standard approach when investigating intercellular communication from scRNA-seq data ^23,24^, although the way of computing the score varies between studies. Here, the choice of the geometric mean (similar to SingleCellSignalR) rather than the arithmetic mean (as used by CellPhoneDB) is motivated by several advantages. The first one is that the geometric mean tends towards zero if either *e*_*L*_ or *e*_*R*_ tends to zero. This implies that when a highly expressed ligand is combined with a lowly expressed receptor (or vice-versa) the score is not dominated by the large ligand value, as it would have been the case with the arithmetic mean. Along the same line, although transcript counts or UMIs only give an indirect representation of protein levels ^92^, molecular interactions are usually modeled by the law of mass action ^93^ which is by essence multiplicative and not additive in protein concentrations. Finally, the geometric mean provides a clear interpretation of the log fold-change of the scores between the two conditions of interest. Indeed, we see that the log fold-change of the CCI score across two hypothetical conditions *A* and *B* corresponds to the (arithmetic) average between the respective ligand and receptor log fold-changes, 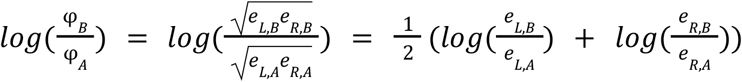.

### CCI detection and differential analyses

Our approach relies on three permutation tests to assess if a CCI is 1) cell-type pair specific in condition *A*, 2) cell-type pair specific in condition *B* and 3) differentially expressed between *A* and *B*. To be computationally more efficient, the three tests are done together as part of a single iteration loop. All threshold parameters described below can be adjusted by the users.

Given *m* cell types and *l* LRIs that are found in the scRNA-seq dataset, scDiffCom builds a table of *m*^2^ · *l* hypothetical CCIs. For each CCI, we compute the CCI scores φ_*A*_ and φ_*B*_ , the log fold change *log*(φ_*B*_ /φ_*A*_) and the variables *n*_*i,j*_ and *d*_*i,j*_ corresponding to the number and fraction of emitter cells expressing the ligand (*i* = *L*) or receiver cells expressing the receptor (*i* = *R*) in either condition (*j* ∈{*A, B*}). A CCI is deemed *not expressed* in condition *j* if (*n*_*L,j*_ < 5 or *n*_*R,j*_ < 5) and (*d*_*L,j*_ < 0. 1 or *d*_*R,j*_ < 0. 1).

Only the CCIs *expressed* in at least *A* or *B* are passed to the iteration loop. At each iteration *k*, three independent operations are done: 1) shuffling the cell-type labels of cells from condition *A* and returning the random score 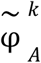 as the *k*-th element of the null distribution representing the random variable Φ_*A*_, 2) same for condition *B*, returning 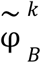 to form the null distribution of Φ_*B*_ , and 3) keeping the original cell-type labels but shuffling the *A* and *B* condition labels, and returning the random score difference 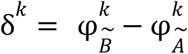 to form the null distribution of the random variable ∆ After iterating, the true values φ_*A*_, φ_*B*_ and δ = φ_*B*_ − φ_*A*_ are compared to the three null distributions in order to compute the two one-sided specificity p-values *p*_*A*_ = *P*(Φ_*A*_ > φ_*A*_) and *p*_*B*_ = *P*(Φ_*B*_ > φ_*B*_), and the differential two-sided p-value *p*_*DE*_ = *P*(|∆| > |δ|). Those p-values are then adjusted for false discovery rate according to the Benjamini-Hochberg procedure ^94^.

A CCI is considered *detected* in condition *j* ∈{*A, B*} if 1) it is *expressed*, 2) it is *specific*, based on the specificity p-values (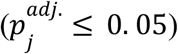, and 3) its score is among the top 80% of all the *specific* CCI scores of both conditions, namely 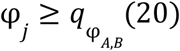, where 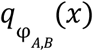 is the x-th percentile of the scores. A CCI is called *differentially expressed* if 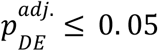 and *differentially expressed* if 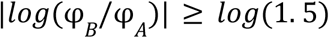.

### CCI classification

We only kept CCIs *detected* in at least one of the two conditions. They are then classified into four categories: 1) *UP* when 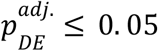 and *logfc* ≥ *log*(1. 5), 2) *DOWN* when 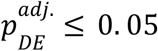 and *logfc* ≤− *log*(1. 5), 3) *FLAT* when |*logfc*| < *log*(1. 5) and 4) *NSC* (non-significant change) when 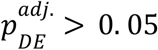 and |*logfc*| ≥ *log*(1. 5).

The detection analysis was used to remove biologically irrelevant interactions, but not to predict actual changes. Using the detection test for this purpose was indeed prone to return false-positive varying signals, i.e., CCIs that seem to appear or disappear because they fluctuate around the detection threshold, but which are in reality not differentially expressed. Using the aging datasets as benchmarking data, we considered all possible outcomes (Supplemental Data 2) and only noticed a marginal number of seemingly contradictory cases between the two tests, such as disappearing CCIs with positive log fold change. Those are for instance due to a reduction of the fraction of expressing cells, despite the increase of the signal, hence our decision to prioritize the classification of the CCIs based on the differential test.

### Fisher’s exact test to find over-represented signals

Over-representation analysis (ORA) is used to evaluate frequent patterns in categorical data, for example, to find if a particular feature of CCIs, e.g., the annotation with the GO term *T cell differentiation*, is more frequent in *up-regulated* CCIs compared to all other CCIs. This statistical association is measured by compiling the corresponding 2×2 contingency table (*up-regulated*/*not up-regulated* vs *annotated*/*not annotated*) and applying Fisher’s exact test. We performed this procedure for every CCI feature (all GO Terms, KEGG pathways, LRIs, ligands, receptors, cell types) and classes (UP, DOWN, FLAT). It returns an odds ratio (OR) and a p-value adjusted for multiple testing according to the Benjamini-Hochberg procedure ^94^. In some instances (e.g., pattern ranking, plots), to sort the results based on a single value, we combined the OR and the p-value to create an ORA score, by adapting the gene-significance score (π-value) used in differential gene expression analysis and GSEA ^95^: *ORA score* = *log*_2_ *OR* · (− *log*_10_ *pval*).

### Visualization Tools in scDiffCom

We implemented two functions in scDiffCom to visualize the over-representation results. scDiffCom::PlotORA displays the top over-represented keywords of a given category and regulation. It is implemented on top of the R package ggplot2 ^96^. scDiffCom::BuildNetwork shows on a summary graph the over-represented cell types and cell-type pairs. It relies on the R package igraph ^97^ for internal computations and on the R package visNetwork ^98^ for the interactive rendering.

### Fetching and preparing scRNA-seq datasets

We downloaded the latest version (as of March 21, 2021) of Tabula Muris Senis from the Amazon S3 czb-tabula-muris-senis repository and the Calico dataset from the calicolabs website. They had been preprocessed and annotated with the python toolkit Scanpy ^99^ prior to our work, and we converted the resulting h5ad files to R Seurat objects.

As stated in the original TMS article ^38^, FACS and Droplet refer to the technique used to capture the cells, namely 1) cell sorting in microtiter well plates followed by Smart-seq2 library preparation and full-length sequencing and 2) cell capture by microfluidic droplets as per the 10x Genomics protocol followed by 3’ end counting. The Calico data were exclusively obtained using the Droplet technique. Regarding the mice age, TMS provided multiple time points that needed to be grouped into *young* and *old* categories. We removed 1-month-old and 30-month-old cells to avoid bias due to developmental or longevity-related processes. Therefore, we compared 3-month-old to 18/24-month-old cells from TMS and 7/8-month-old cells to 22/23-month-old cells from Calico. Finally, we filtered out tissues that were missing one age group (e.g., *TMS Droplet Fat* only contained old cells).

The cells from each dataset were sequenced together. However, we decided to regroup *TMS FACS Brain_Myeloid* and *Brain_Non-myeloid* as the former only contained two cell types (*macrophage* and *microglial cell*) and that merging the datasets allowed us to infer interactions with the other parts of the brain. We verified that this did not significantly alter the interactions detected in each dataset independently.

### Cell-type characterization

In each dataset, we standardized the names of the cell types based on Cell Ontology standards ^100^, e.g., *atrial myocyte* was renamed as *regular atrial cardiac myocyte*. We also regrouped some specialized cell clusters, e.g., CD4+ and CD8+ T cells, in order to increase sample size and avoid overlapping cell types. These overlaps were exceptionally kept in some tissues, for instance, the *undetermined myeloid leukocytes* in the Lung dataset from *Calico Droplet (male)* overlap with some specialized cell types such as *classical monocytes*, but were worth keeping as distinct categories. Finally, we classified the cell types in 10 families to facilitate downstream analyses: *endothelial cells, epithelial cells, connective tissue cells, leukocytes, stem cells, neurons, glial cells, muscle cells, erythroid lineage cells*, and *hematopoietic precursor cells*. The list of all cell types with their new names and family annotations is available in Supplemental Data 3.

### Building and deploying scAgeComShiny

We used the R package golem ^101^ to build the Shiny App scAgeComShiny which contains all scAgeCom results. Interactive scatter plots were built with plotly ^102^, which was also used to display the GO Terms treemaps. Those were internally computed with the R package rrvgo ^103^, according to the original method from REVIGO ^104^. To deploy the application, we first used golem to create a Docker image of the scAgeComShiny app and then serve it with the containerized version of ShinyProxy open source middleware.

## Acknowledgements

We thank Jacob Kimmel for helpful information regarding the Calico murine cell atlas, Koki Tsuyuzaki for general discussion regarding intercellular communication at the early stage of this project, Zoya Abidi for her support to deploy the Shiny app and Ulysse Castet for his useful suggestions on the manuscript. Final figures were created with BioRender.com. CL is grateful for the funding provided by the Human Frontier Science Program (fellowship LT000741/2019-C). Our work in the Integrative Genomics of Ageing Group is further supported by grants from the Wellcome Trust (208375/Z/17/Z), LongeCity and the Biotechnology and Biological Sciences Research Council (BB/R014949/1) to JPM. RT and EU are supported by the National Authority for Scientific Research and Innovation, and by the Ministry of European Funds, Romania, through the Competitiveness Operational Programme 2014-2020, POC-A.1-A.1.1.4-E-2015 [Grant number: 40/02.09.2016, ID: P_37_778, to RT].

## Author contributions

CL: conceived the project, developed scDiffCom/scAgeCom/scAgeComShiny, compiled the ligand-receptor database, preprocessed scRNA-seq datasets, interpreted the results, wrote the manuscript

EU: developed scDiffCom/scAgeCom/scAgeComShiny, curated the ligand-receptor database, interpreted the results, wrote the manuscript

AE: contributed to scDiffCom/scAgeComShiny, annotated the ligand-receptor database, annotated the TMS/Calico cell types, interpreted the results, wrote the manuscript

RA: preprocessed scRNA-seq datasets, interpreted the results, revised the manuscript

AP: annotated the TMS/Calico cell types

RT: revised the manuscript, supervised the project

JPM: revised the manuscript, supervised the project

## Code availability

- R package scDiffCom: https://github.com/CyrilLagger/scDiffCom
- R scripts for the aging analysis: https://github.com/CyrilLagger/scAgeCom
- Golem package for the Shiny app: https://github.com/CyrilLagger/scAgeComShiny
- Docker image of the Shiny app: https://hub.docker.com/r/ursueugen/scagecom
- scAgeCom website: http://scagecom.org/

